# Dopaminergic Medication Improves Force Control and Modifies Motor Unit Activity During the Steadiest and Least Steady Periods in Parkinson’s Disease Force control and motor units in Parkinson’s Disease

**DOI:** 10.64898/2026.10.16.758033

**Authors:** Parisa Alaei, Aleš Holobar, Daryl J. Wile, Jennifer M. Jakobi

**Author notes:** Correspondence: Jennifer M. Jakobi, PhD The University of British Columbia Okanagan 150-1147 Research Road (Arts Building) Kelowna, BC Canada V1V 1V7. Authors emails: Parisa Alaei Aleš Holobar Daryl J. Wile Jennifer M. Jakobi.

## Abstract

Parkinson’s disease (PD) impairs steady force production, but how dopaminergic medication alters force control and the motor unit (MU) activity is unclear. This study examined force steadiness and biceps brachii MU behavior in 16 people with PD ON and OFF dopaminergic medication. Participants performed sustained isometric elbow flexion contractions at 5%, 10%, 15%, and 25% of maximum voluntary contraction (MVC). High-density surface electromyography was decomposed into MU spike trains. Force steadiness and MU discharge properties were quantified during nonoverlapping 7-s steadiest and least steady contraction periods, whereas MU coherence was assessed across the force plateau. In the ON state, force steadiness improved by ∼17–19% during both contraction periods at 5–15% MVC. MU interspike interval variability decreased by ∼12% across all forces during the least steady contraction period and by ∼17% at 15% MVC during the steadiest contraction period, whereas mean MU discharge rate was unchanged. Recruitment thresholds were higher at 10% and 25% MVC in the ON state, whereas derecruitment thresholds did not differ. Across the entire plateau, alpha- and tremor-band MU coherence decreased by ∼14% and ∼18%, respectively, whereas delta-band coherence was unchanged. These findings demonstrate that dopaminergic medication improves both the steadiest and least steady periods of low-force control, accompanied by reduced MU discharge variability and frequency-specific changes in plateau-wide MU coherence rather than changes in mean discharge rate. Resolving within-contraction variation reveals medication-sensitive features of force control that are obscured by conventional plateau-wide analyses, providing a new approach for evaluating motor output in PD.

## Significance Statement

Parkinson’s disease disrupts the ability to maintain steady muscle force, limiting precise control of movement. We show that dopaminergic medication improves force steadiness during both the steadiest and least steady periods of sustained low-force contractions. Medication also reduced motor-unit discharge variability most consistently during the least steady periods and altered alpha- and tremor-band coherence across the motor-unit pool without changing mean discharge rate. These findings indicate that dopaminergic medication may enhance the neural regulation of force. By resolving the best and poorest periods of low-force control within individual contractions, this study identifies medication-sensitive changes in motor-unit activity and force output that conventional analyses of the entire contraction or only its steadiest portion may obscure.

## Introduction

Precise regulation of muscle force is essential for motor function and is impaired in Parkinson’s disease (PD). Degeneration of dopaminergic neurons in the substantia nigra pars compacta alters basal ganglia circuitry and produces tremor, bradykinesia, rigidity, and postural instability (Dauer & Przedborski, 2003; Jankovic, 2008). Impaired motor control is also evident in the reduced ability to maintain steady force during submaximal isometric contractions (Brown, 1997; Ko et al., 2015; Smart et al., 2020). Force steadiness reflects both variability in the discharge of individual motor units (MUs) and common synaptic input to the MU pool (Enoka & Farina, 2021). How dopaminergic medication modifies these features of MU activity and their expression in force output remains unclear.

Studies of individual MUs in PD have identified lower mean discharge rates, primarily in intrinsic hand muscles (Dietz et al., 1974; Milner-Brown et al., 1979), greater discharge variability (Kim et al., 2024; Wilson et al., 2020), and lower recruitment thresholds in the first dorsal interosseous (Glendinning & Enoka, 1994). Greater estimated motoneuron excitability has also been reported in PD (Alaei et al., 2026), and associations between striatal dopamine binding and MU discharge characteristics link dopaminergic dysfunction to MU behavior (Nishikawa et al., 2021). However, most previous studies compared people with PD and neurologically intact adults or examined MU behavior in only one medication state. Consequently, it remains unclear how individual MU discharge properties change when the same individuals are assessed during the ON and OFF dopaminergic states. It is also unknown whether medication alters common input across the MU pool. Coherence between cumulative MU spike trains provides a frequency-specific estimate of this common input and complements measures of individual MU discharge (Farina et al., 2014; Negro & Farina, 2011).

Force steadiness has typically been quantified across the entire force plateau (Hortobágyi et al., 2001; Kim et al., 2024) or during its steadiest segment (Daneshgar et al., 2024; Tvrdy et al., 2025). Plateau-wide analyses characterize overall force control, whereas selection of the steadiest segment characterizes the best control achieved during the contraction. Neither approach captures the range of force variability that can occur within a sustained contraction. Comparing the steadiest and least steady periods provides a means of determining whether individual MU discharge properties differ between periods of better and poorer force control and whether dopaminergic medication modifies these differences. In contrast, MU coherence characterizes the frequency-specific organization of MU-pool activity and addresses the separate question of whether this overall organization differs between medication states.

This study aimed to determine how dopaminergic medication influences force control and MU behavior during sustained submaximal isometric contractions in the more affected upper limb of people with PD. Force steadiness and individual MU discharge properties were compared between the steadiest and least steady contraction periods, while MU coherence was evaluated across the entire force plateau. We hypothesized that dopaminergic medication would improve force steadiness during both periods and that this improvement would be accompanied by less variability in MU discharge. We further hypothesized that medication would modify the coordination of MU activity across the contraction.

## Methods

### Participants

Eleven males and five females with mild to moderate PD severity participated in this study. All participants had a confirmed diagnosis of PD and were undergoing treatment with neurologist-prescribed levodopa-carbidopa. They completed two identical sessions in a randomized order between ON and OFF medication states. Participants were assessed in the OFF-medication state after withholding dopaminergic medication for at least 12 hours and in the ON medication state, 60–90 minutes after medication intake typically in the morning (6 of 16 ON states occurred in afternoon for patient feasibility). Clinical assessments were performed first in each medication state, followed by the experimental force-control and motor-unit assessments. The Hoehn and Yahr rating was assessed exclusively during the OFF state, whereas motor symptom severity during both the ON and OFF states was assessed using the total MDS-UPDRS III score and the MDS-UPDRS III score for the participant’s self-reported more affected upper limb. Individuals with cardiac conditions, any recent major injuries, or surgeries to the more affected limb, which was being tested, within the last six months, or neurological disorders other than PD were excluded. This study was approved through the harmonized review process (H22-02896) by the Clinical Research Ethics Board of the University of British Columbia and Interior Health and was conducted in accordance with the Declaration of Helsinki. All participants provided written informed consent prior to starting the study.

### Experimental Set-up and Procedure

Participants underwent neuromuscular testing while seated in a custom-built elbow flexion dynamometer. The setup was adjusted to ensure that participants feet were supported on a flat surface, their knees and hips were flexed at 90°, the shoulder was slightly abducted (∼15°), and the elbow was positioned at 110° of flexion. Participants gripped the manipulandum with the hand on their self-reported more affected side, while their forearm was supported on a padded rest in a neutral position. To minimize any extraneous movement, a non-distensible strap was used to secure them to the chair (Figure 1). Isometric elbow flexion force was sampled at 1004 Hz using a Power 1401 data acquisition system (Cambridge Electronic Design (CED), Cambridge, England), amplified 100x, and stored for offline analysis in Spike2 (v10.04 software, CED, Cambridge, England).

**Figure 1.**
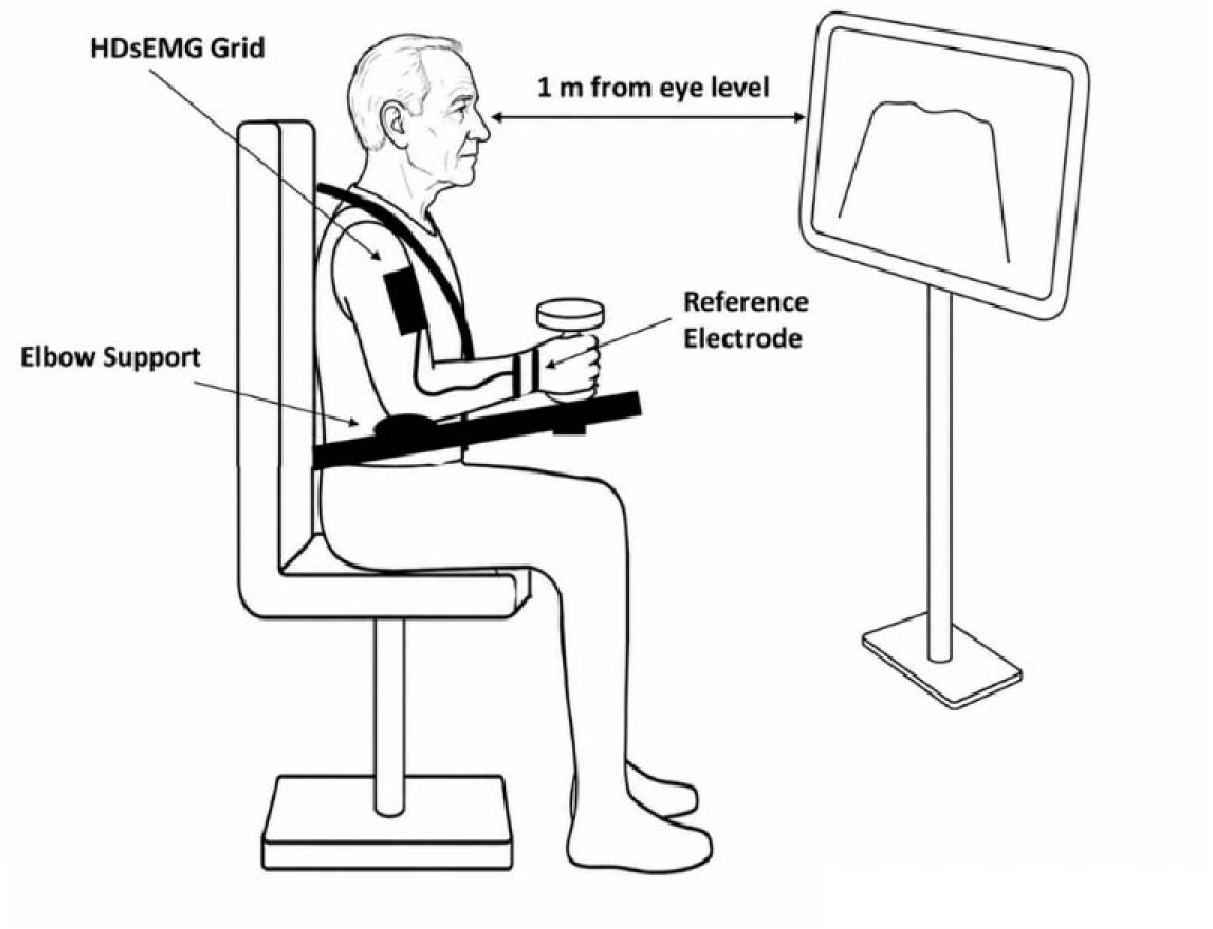
Representative diagram of the experimental set-up. Participants sat in a custom-built elbow flexion dynamometer chair with hips and knees flexed at 90° and the elbow at 110°. Participants gripped a manipulandum on the more affected side in a neutral forearm orientation. The force measurements were displayed in real-time on a 52 cm monitor positioned 100 cm in front of the participant with the center of the screen at eye level. The base image was generated using ChatGPT and modified by the authors to accurately represent the experimental setup

### Maximum Voluntary Contraction and Submaximal Tracking Tasks

To assess maximum voluntary contraction (MVC), participants were instructed to pull upward against the chair manipulandum as hard and as fast as possible. Following a brief practice period, participants performed three to five isometric elbow flexion MVCs, each held for 3 to 5 seconds. Approximately 2 minutes of rest were provided between contractions.

Relative submaximal force levels of 5, 10, 15, and 25% MVC were used for the submaximal force tracking tasks and calculated based on the highest MVC attained at each session. Participants then performed isometric tracking tasks in the neutral position, with the order of force levels randomized. The tracking task involved a 3-second resting state, followed by a gradual increase in force to the target level at a rate of 10% MVC per second. This was followed by a 25-second steady hold at the target force before gradually returning to baseline at the same rate. Before the experimental trials, participants completed 1–2 practice trials to become familiar with the force-tracking task. Submaximal tracking tasks were performed three to four times at each force level. The visual feedback was scaled relative to each participant’s MVC and all participants received identical feedback by scaling different absolute forces to appear in an identical location and sensitivity on the screen.

### High-density surface EMG

Before electrode placement, the skin regions for the grid electrodes and reference electrodes were lightly abraded and cleaned with alcohol pads to reduce impedance and improve signal quality. High-density surface EMG (HDsEMG) electrodes (64 electrodes, 13x5, 8mm I.E.D., GR08MM1305, OT 126 Bioelettronica, Inc., Turin, IT) were placed over the long axis of the biceps brachii using double-sided adhesive foam (FOA08MM1305, Bioelettronica, Inc., Turin, IT) with conductive gel used to minimize electrode impedance. A reference electrode strap was placed around the wrist. HDsEMG data was recorded in single differential mode, at 2048 Hz, and band-pass filtered at 10-500Hz using a Quattrocento amplifier (OT Bioelettronica, Inc., Turin, IT). HDsEMG and force signals were synchronized with Transistor-Transistor Logic (TTL) pulse at the onset of each trial.

### Data Analysis

All recorded data (force and HDsEMG) were converted offline to MATLAB (MATLAB R2024a; MathWorks Inc., Natick, MA, USA) for further processing. The force signal was low-pass filtered at 30 Hz using a fourth-order Butterworth filter. Force steadiness was analyzed and is reported for the steadiest and least steady 7-second periods of the 25-s plateau state. Using automated MATLAB code to calculate the SD of force, the steadiest and least steady non-overlapping 7-s periods were determined, excluding the first and last 2 s to minimize the influence of contraction onset and termination. Force steadiness was calculated as the coefficient of variation (CV) of force, defined as: CV = (standard deviation of force / mean force) x 100. To determine whether the least and most steady periods occurred at the beginning or end of the contraction, we identified the portion of the contraction in which 50% or more of that period fell within either the first or last half of the contraction.

Monopolar HDsEMG signals were initially bandpass filtered (10−500 Hz), and all channels exhibiting low correlation with their direct neighbors were removed (typically <5% of all channels). Convolutive blind source separation was used to separate HDsEMG recordings into individual MU discharges (Holobar and Zazula 2007). The MU discharge trains were visually inspected and manually edited in DEMUSE software (v. 6.0; the University of Maribor, Slovenia). Data were excluded from the analysis when fewer than 10 consecutive discharges were identified or when interspike intervals (ISIs) were <33.3 ms, corresponding to a firing rate >30 Hz. Across all included MUs, the average PNR was 30.8 ± 4.6 dB. For each MU spike train, the average MUDR (Hz) and the coefficient of variation of interspike intervals (CVISI, %) were calculated for the periods that corresponded to the steadiest and least steady 7-second period of the plateau state. The recruitment (%) and derecruitment (%) thresholds were defined as the instantaneous force values of the first and last spike in the series of action potentials, respectively, and reported as a relative force (% MVC).

Coherence analysis was performed over the entire plateau state of contraction in both the ON and OFF medication states to quantify frequency-dependent synchronization between MU discharge patterns. To provide sufficient temporal support for reliable coherence estimation, the entire plateau phase was used for the coherence calculation. MUs were randomly divided into two independent groups containing approximately equal numbers of MUs. For each trial, this randomization procedure was repeated 100 times to minimize the influence of MU selection. For each MU group, a normalized cumulative spike train (CST) was generated by summing MU discharge events and normalizing by the number of active MUs. Coherence was then calculated between the two normalized CSTs using the method described by Raethjen et al. (2007). Coherence was estimated using a 0.75-s Hann window with no overlap between consecutive segments, a frequency resolution of 0.25 Hz, and within a frequency range of 0–100 Hz. Coherence values were Fisher Z-transformed using: 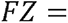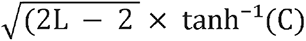 where C represents the coherence value, and L represents the number of analyzed epochs. The 95% confidence limit for significant coherence was calculated based on the number of analyzed epochs. Coherence spectra from the 100 random MU subdivisions were averaged to obtain the final coherence spectrum for each trial. Mean coherence values were calculated within predefined frequency bands: delta (0–5 Hz), tremor (3–12 Hz), alpha (5–15 Hz), beta (15–30 Hz), and gamma (30–60 Hz)(Cabral et al., 2024; Reithe et al., 2026; Vanmechelen et al., 2023; Yu et al., 2025).

### Statistical Analysis

R (version 4.3.2; R Foundation for Statistical Computing) was used for all analyses. To characterize participants and examine differences in force steadiness and MU properties, linear mixed-effects (LME) models were employed. The primary model was defined by medication state (ON vs. OFF) and contraction steadiness (OFF least steady, ON least steady, OFF steady, and ON steady). The OFF least steady condition was treated as the reference (least ideal condition), and all other conditions were compared against it. Dependent variables included force steadiness, CVISI, and MUDR. Post hoc comparisons among conditions were performed using estimated marginal means.

For recruitment threshold and derecruitment threshold, the LME model structure included the four levels of, OFF recruitment threshold, ON recruitment threshold, OFF derecruitment threshold, and ON derecruitment threshold, with OFF recruitment threshold set as the reference level. Model residuals were checked for normality and homoscedasticity. As the residuals for CVISI, CV of Force, Recruitment and derecruitment did not meet the normality assumption, logarithmic transformations were applied to these variables before analysis. Recruitment and derecruitment thresholds were not normally distributed, and thus assessed with Spearman’s rank correlation

For coherence, the primary linear mixed-effects model included Force (5%, 10%, 15%, and 25% MVC) and medication state (ON vs. OFF) as fixed effects, as well as their interaction (Force × Medication). Participants were included as a random intercept to account for repeated observations within participants. The dependent variable was Fisher z-transformed coherence, and analyses were performed separately for each frequency band (delta, alpha, beta, gamma, and tremor). Type III ANOVA with Satterthwaite’s method was used to test the main effect of the fixed factors. Post hoc comparisons were performed using estimated marginal means (emmeans).

For comparisons of MDS-UPDRS III scores and MVC between ON and OFF states, paired-sample t-tests were used. Pearson’s correlation was used to examine relationships between affected upper limb MDS-UPDRS III scores and CV of force, MUDR, and CVISI. The probability level for all statistical tests was set at 0.05, and results are reported as mean ± SD in the text and figures. LME models were fit using the lme4 package (Bates et al., 2015), with p-values estimated via Satterthwaite’s method (lmerTest package; Kuznetsova et al., 2017).

## Results

Eleven males (68 ± 8.1 yrs; 179.1 ± 11.1 cm; 86.0 ± 13.5 kg) and five females (72.6 ± 7.0 yrs; 163.2 ± 7.3 cm; 57.6 ± 6.6 kg) with mild to moderate PD severity (Hoehn Yahr Score OFF state, 1.6 ± 0.6; range 1-3; PD Diagnosis Duration 6.4 ± 5.8 years; Dopa equivalent, 487.5 ± 294.7 mg/day) participated in this study. The total MDS-UPDRS III (OFF: 23.2 ± 9.5, ON: 16.9 ± 9.5; t_15_ = 5.6, p < 0.001) and the MDS-UPDRS III for the more affected upper limb (OFF: 5.7 ± 2.0, ON: 4.3 ± 1.9; t_15_ = 3.0, p = 0.01) was significantly lower during the ON state compared with the OFF state of medication. During the ON state (177.5 ± 60.2 N) compared with OFF (170 ± 57.5 N), elbow flexor MVC was significantly higher (t_15_ = 2.4, p = 0.03).

### Motor Unit Properties

Decomposition from the biceps brachii of the affected side in sixteen people with PD during submaximal isometric elbow flexion contraction yielded a total of 1485 MU spike trains (OFF: 737, ON:748) during the least steady and 1469 MU spike trains (OFF: 727, ON:742) during the steadiest periods across all forces (Table 1; Figure 2). Of these MUs, 1423 were recorded in both the least steady and steadiest periods.

**Figure 2.**
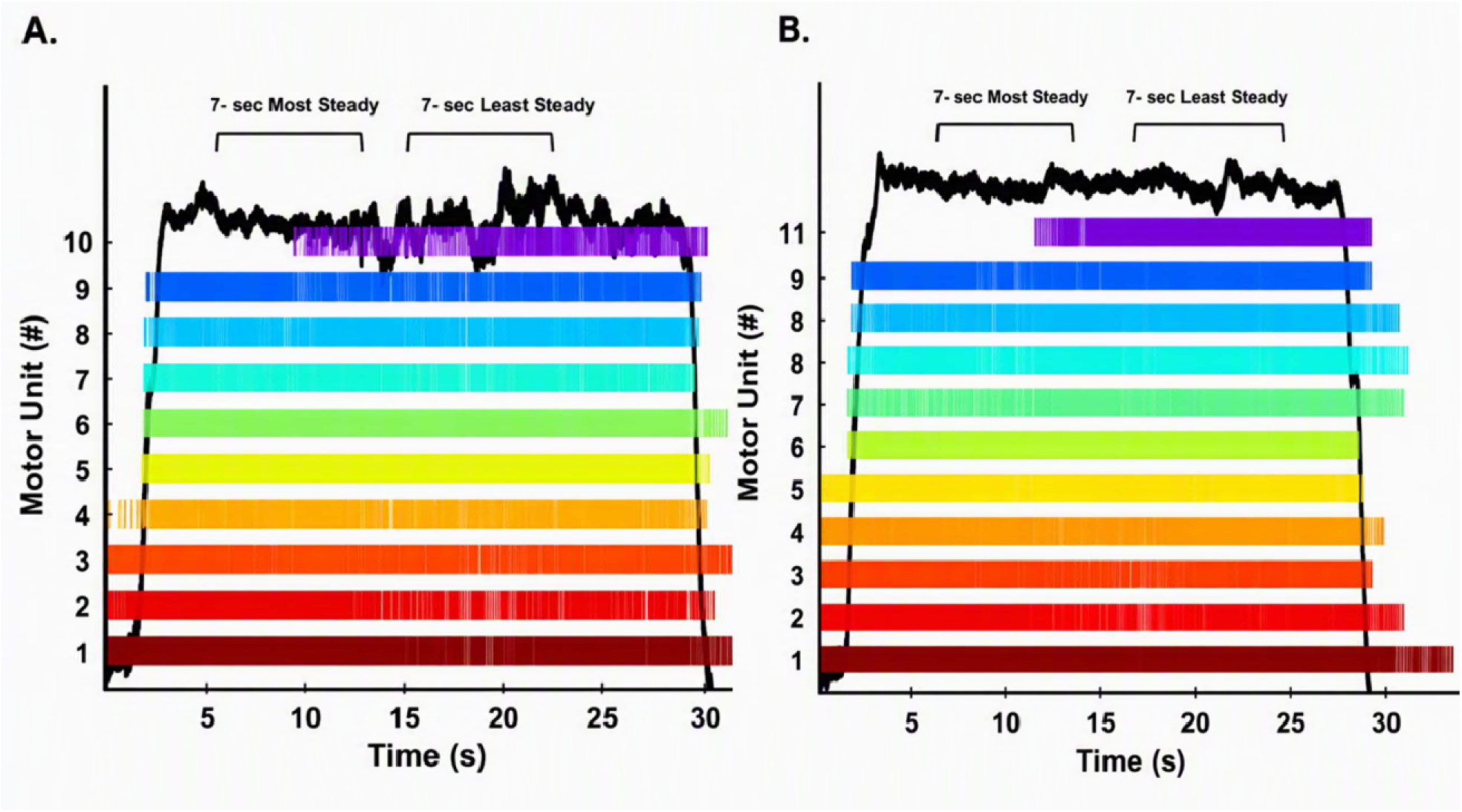
Representative Data of the force tracking task (Black Line both panels) at 10% MVC for a male during the (A) OFF state with discharge trains of 10 MUs and (B) ON state with discharge trains of 11 motor units. The 7-second most steady and least steady epochs are indicated using brackets. s, second. #, Number of motor units.

**Table 1.** Distribution of recorded motor unit spike trains by condition and force level in ON and OFF states.

| Force Level | Least Steady Period<br>(N = 16; F = 5, M = 11) |  |  |  |  |  | Steady Period<br>(N = 16; F = 5, M = 11) |  |  |  |  |  |
| --- | --- | --- | --- | --- | --- | --- | --- | --- | --- | --- | --- | --- |
|  | OFF |  | ON |  | Total |  | OFF |  | ON |  | Total |  |
|  | F | M | F | M | F | M | F | M | F | M | F | M |
| 5% | 46 | 121 | 46 | 137 | 92 | 258 | 46 | 123 | 48 | 133 | 94 | 256 |
| 10% | 58 | 140 | 57 | 156 | 115 | 296 | 54 | 139 | 57 | 159 | 111 | 298 |
| 15% | 56 | 142 | 56 | 133 | 112 | 275 | 55 | 150 | 55 | 129 | 110 | 279 |
| 25% | 59 | 115 | 49 | 114 | 108 | 229 | 57 | 103 | 45 | 116 | 102 | 219 |
| *Total* | 219 | 518 | 208 | 540 | 427 | 1058 | 212 | 515 | 205 | 537 | 417 | 1052 |

*OFF, OFF state; ON, ON state; N, number; F, number of Females; M, number of Males*

Recruitment threshold was significantly higher at 10% MVC (p = 0.02) and 25% MVC (p = 0.02) during ON compared to the OFF state (see supplemental Table 1 for detailed means and SD). Derecruitment threshold did not change between ON and OFF states at all force levels (p > 0.05). In the OFF state, recruitment threshold was lower than derecruitment threshold at 5% (p < 0.001) and 10% (p = p < 0.001). In the ON state recruitment threshold was lower than derecruitment threshold only at 5% (p < 0.001). Effect-size estimates for these pairwise comparisons are provided in Supplementary Table 2, and the corresponding estimated marginal means, standard errors, and 95% confidence intervals are provided in Supplementary Table 3. Correlation analysis revealed that the MU recruitment thresholds were positively correlated with the derecruitment thresholds across all force levels in both ON (r = 0.54, p < 0.001) and OFF states (r = 0.46, p < 0.001) (Figure 3).

**Figure 3.**
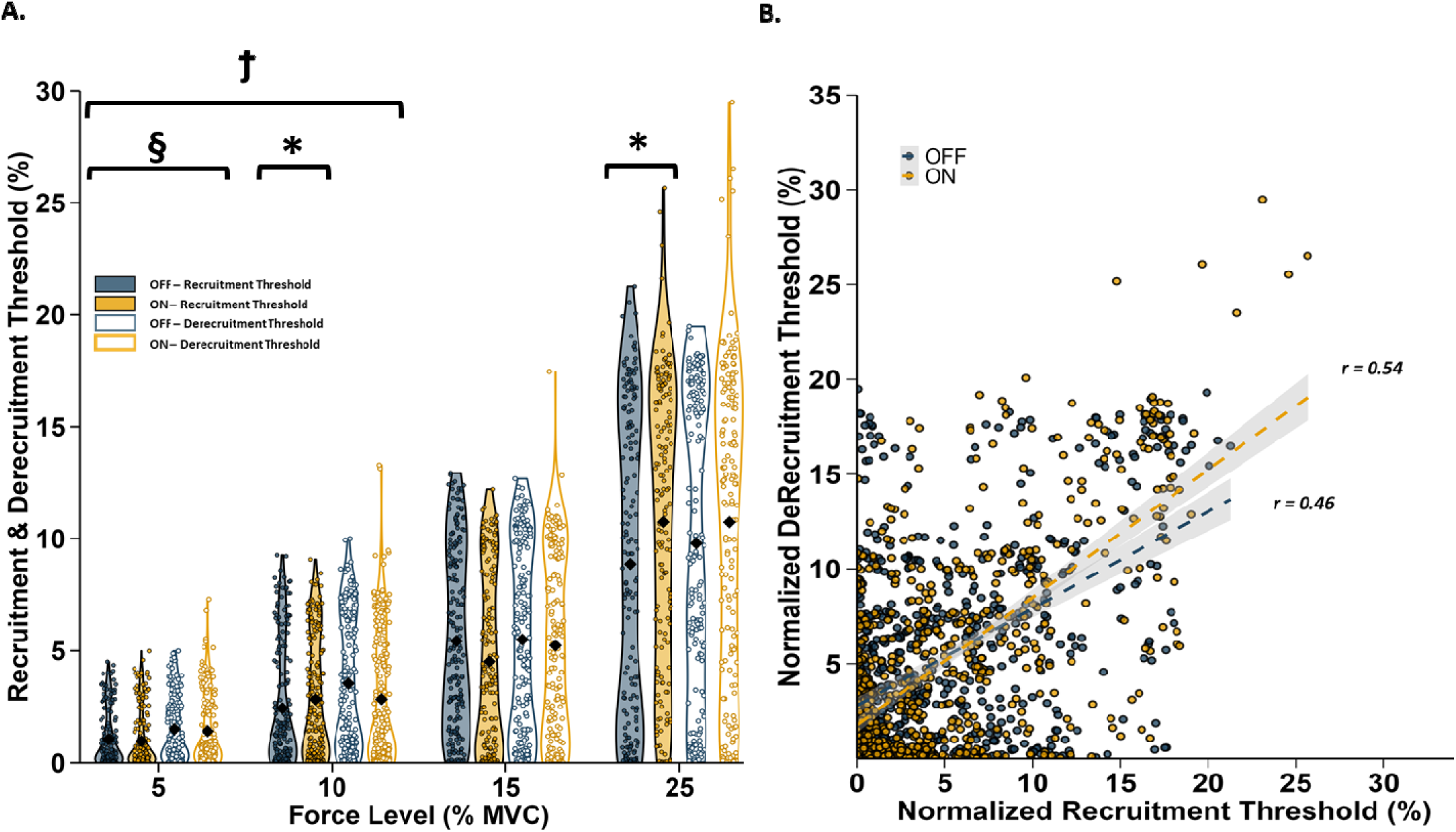
(A) Recruitment and derecruitment threshold box plots in different force levels. (B) Recruitment and derecruitment threshold correlation across all force levels. *, ON differs from OFF state within the Recruitment threshold (P < 0.05); 9, OFF-recruitment threshold differs from OFF-derecruitment threshold (P < 0.05); §, ON-recruitment threshold differs from ON-derecruitment threshold (P < 0.05); Black diamonds represent mean values; Each circle represents an individual data point. r, correlation value.

In the least steady period, MUDR did not differ significantly between the ON and OFF states at any force level (p > 0.05; Figure 4), whereas CVISI was significantly reduced across all force levels during the ON state compared to the OFF state (p < 0.05; Figure 5).

**Figure 4.**
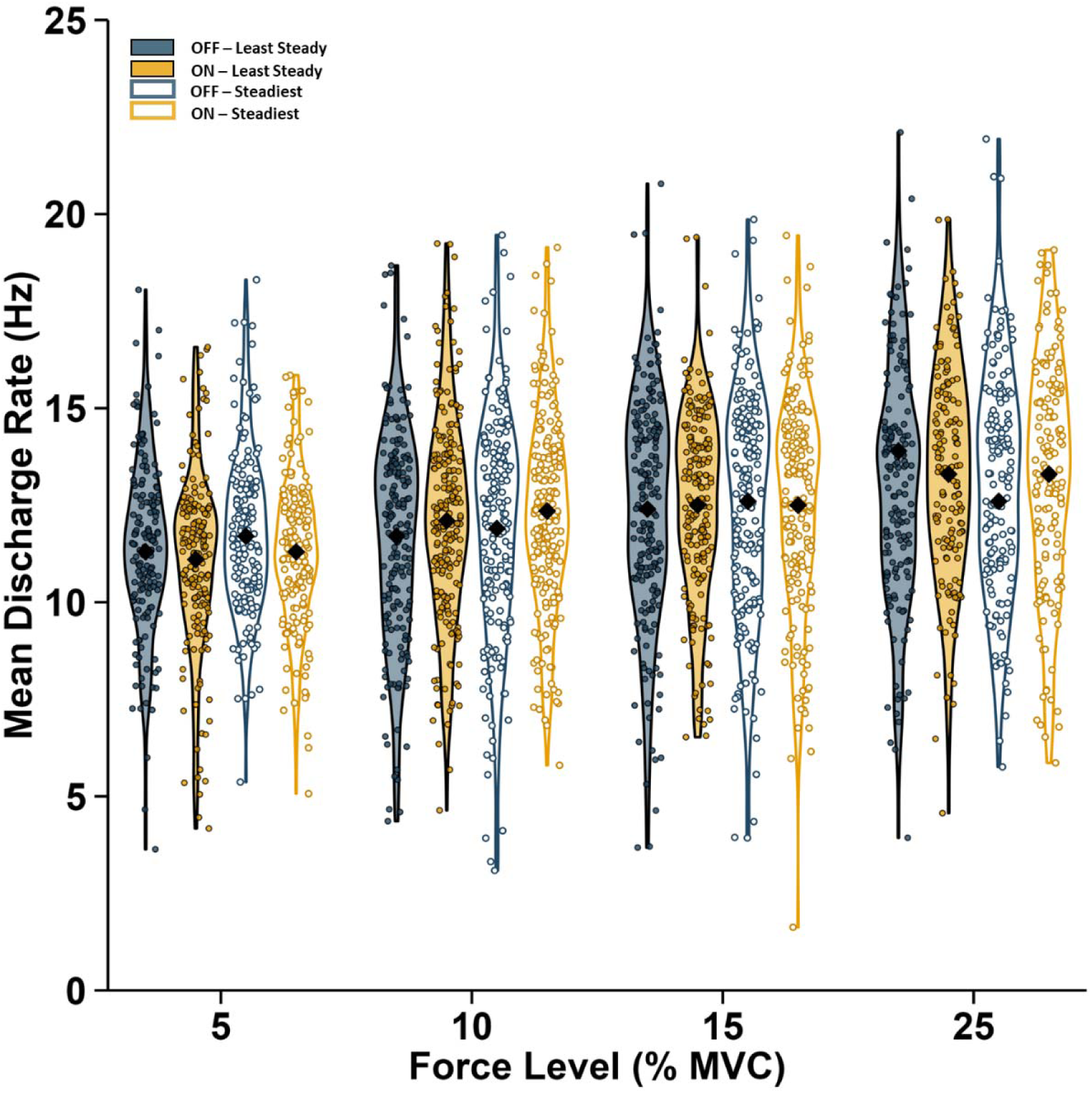
Mean firing rate in the steadiest and least steady periods of an isometric contraction at different force levels. Black diamonds represent mean values; Each circle represents an individual data point; MVC, maximum voluntary contraction; Hz, hertz.

**Figure 5.**
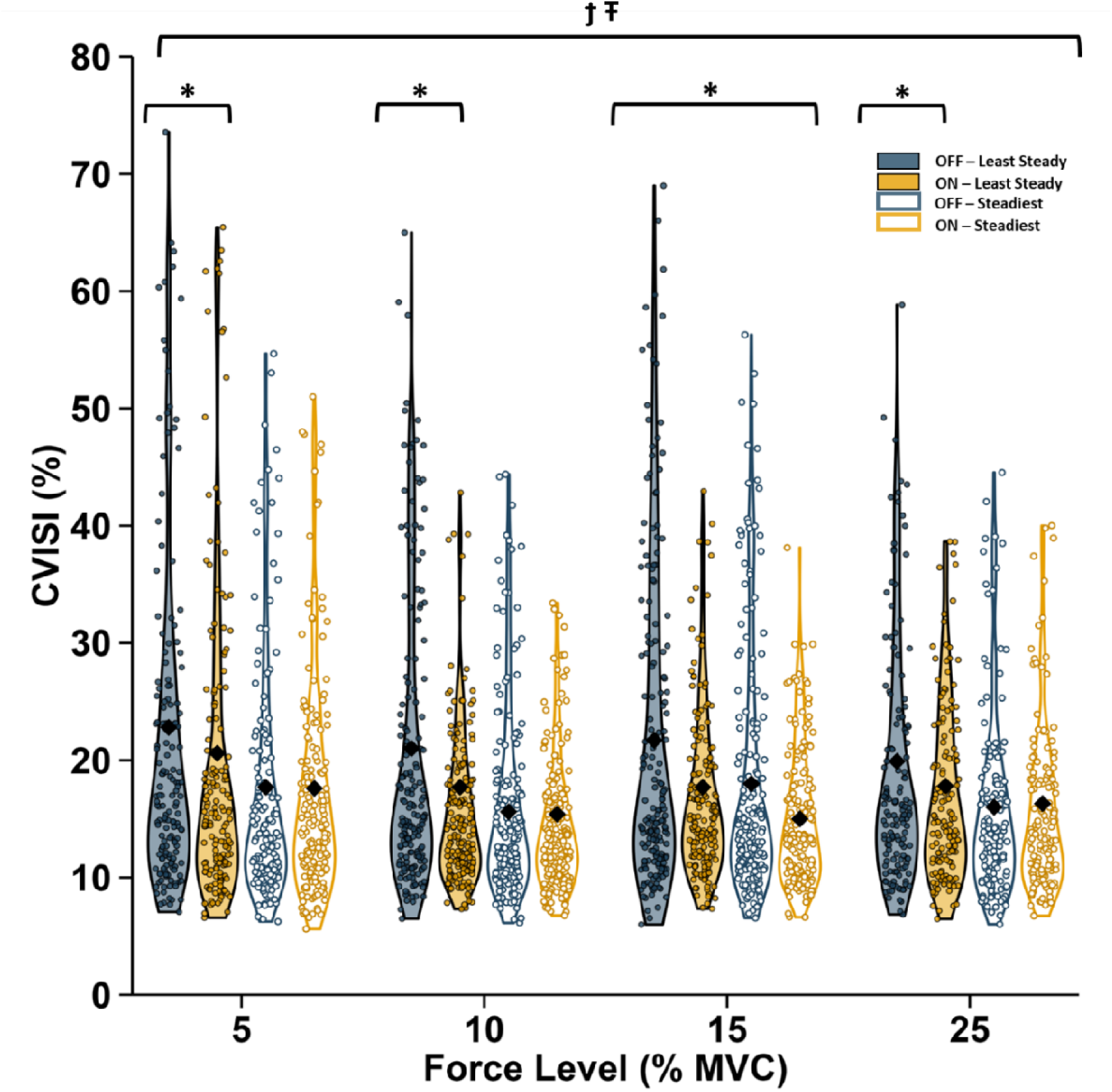
Coefficient of variation of interspike intervals in the steadiest and least steady period of an isometric contraction at different force levels. *, ON differs from OFF states within the period of the least-steady (P < 0.05); 9, OFF least steady differs from ON steadiest across all forces; Ŧ, OFF-least steady differs from OFF-steadiest across all forces. Black diamonds represent mean values; Each circle represents an individual data point; CVISI, Coefficient of variation of interspike intervals; MVC, maximum voluntary contraction.

In the steadiest period, MUDR remained unchanged at all force levels during the ON state compared to the OFF state (p > 0.05; Figure 4). However, CVISI was reduced at 15% MVC (p = 0.03), with no significant differences at other force levels during the ON state compared to the OFF state (p > 0.05; Figure 5).

MUDR did not differ between the steadiest and least steady periods at any force level during either the OFF or ON state (p > 0.05; Figure 4). CVISI was significantly lower during the steadiest period compared to the least steady period across all force levels in the OFF state (p;<;0.05).

However, no significant differences during the ON state were observed in CVISI between the steadiest and least steady periods at any force level (p;>;0.05; Figure 5). Descriptive values for MUDR and CVISI are provided in Supplementary Table 4. Cohen’s d effect sizes for pairwise comparisons of MUDR and CVISI are provided in Supplementary Table 5, and the corresponding estimated marginal means, standard errors, and 95% confidence limits are provided in Supplementary Table 6.

### Force Steadiness

During the least steady period, force steadiness significantly improved in the ON state at 5% (p = 0.02), 10% (p = 0.04), and 15% (p < 0.001) compared to the OFF state (Figure 6). Similar improvements were observed during the steadiest period, with force steadiness significantly enhanced in the ON state at 5% (p = 0.002), 10% (p = 0.03), and 15% (p = 0.01) compared to OFF state (Figure 6). In the OFF state, 56% of the least steady area of the contractions occurred during the second half of the contraction, and similarly in the ON state 53% of the least steady contractions also occurred in the second half of the contraction. Across both medication states, the least steady periods were consistently less steady than the steadiest periods at all force levels. Overall, force control was steadiest during the steadiest period in the ON state compared to the least steady period in the OFF state (p < 0.001; Figure 6). Effect-size estimates for these pairwise comparisons are provided in Supplementary Table 7, and the corresponding estimated marginal means, standard errors, and 95% confidence intervals are provided in Supplementary Table 8.

**Figure 6.**
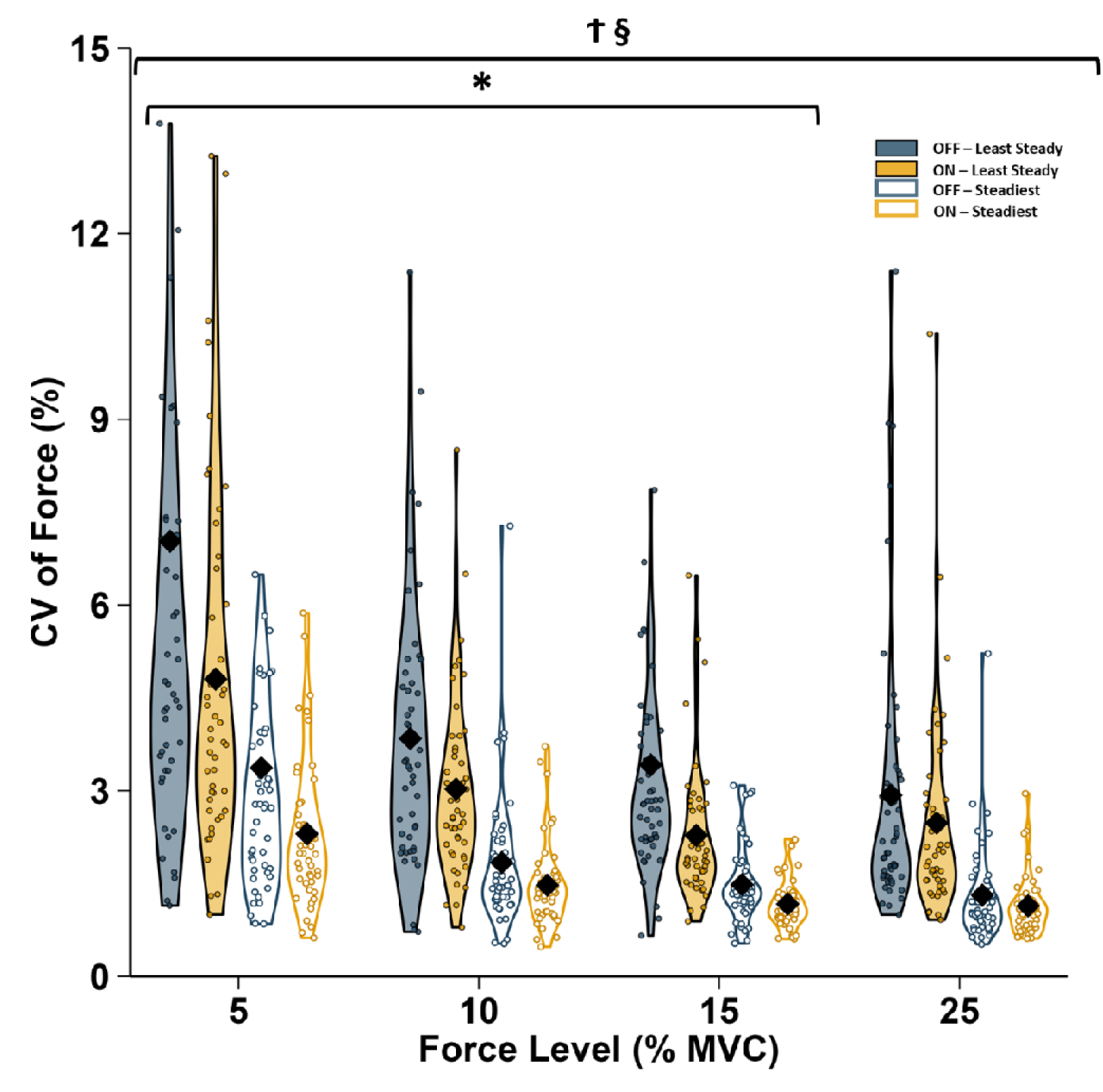
CV of force during the least steady (filled) and steadiest (no-fill) periods of the force signal. *, ON differs from OFF states within the period of the least or steadiest (P < 0.05); §, steadiest period differs from least steady period within the ON and OFF states across all force (P < 0.05); 9, OFF least steady differs from ON steadiest across all forces. Black diamond represent mean values; Each circle represents an individual data point; CV, Coefficient of variation

### MDS-UPDRS III Correlation with Motor Unit and Force Steadiness

The MUDR showed no significant correlation with the affected-side MDS-UPDRS III scores (OFF: R = –0.2, p = 0.4; ON: R = –0.3, p = 0.3) during either the least steady or steadiest periods in both medication states (Figures 7A and 7B). CVISI was also not significantly correlated with MDS-UPDRS III scores (OFF: R = 0.4, p = 0.1; ON: R = –0.1, p = 0.6) in the least steady or steadiest periods (Figures 7C and 7D). Force steadiness did not correlate with MDS-UPDRS III scores during the least steady period in either the OFF (R = 0.3, p = 0.2) or ON (R = 0.01, p = 1.0) medication states (Figure 7E). However, in the steadiest period, a moderate positive correlation was observed in the OFF state (R = 0.5, p = 0.03), while no correlation was found in the ON state (R = 0.1, p = 0.7; Figure 7F).

**Figure 7.**
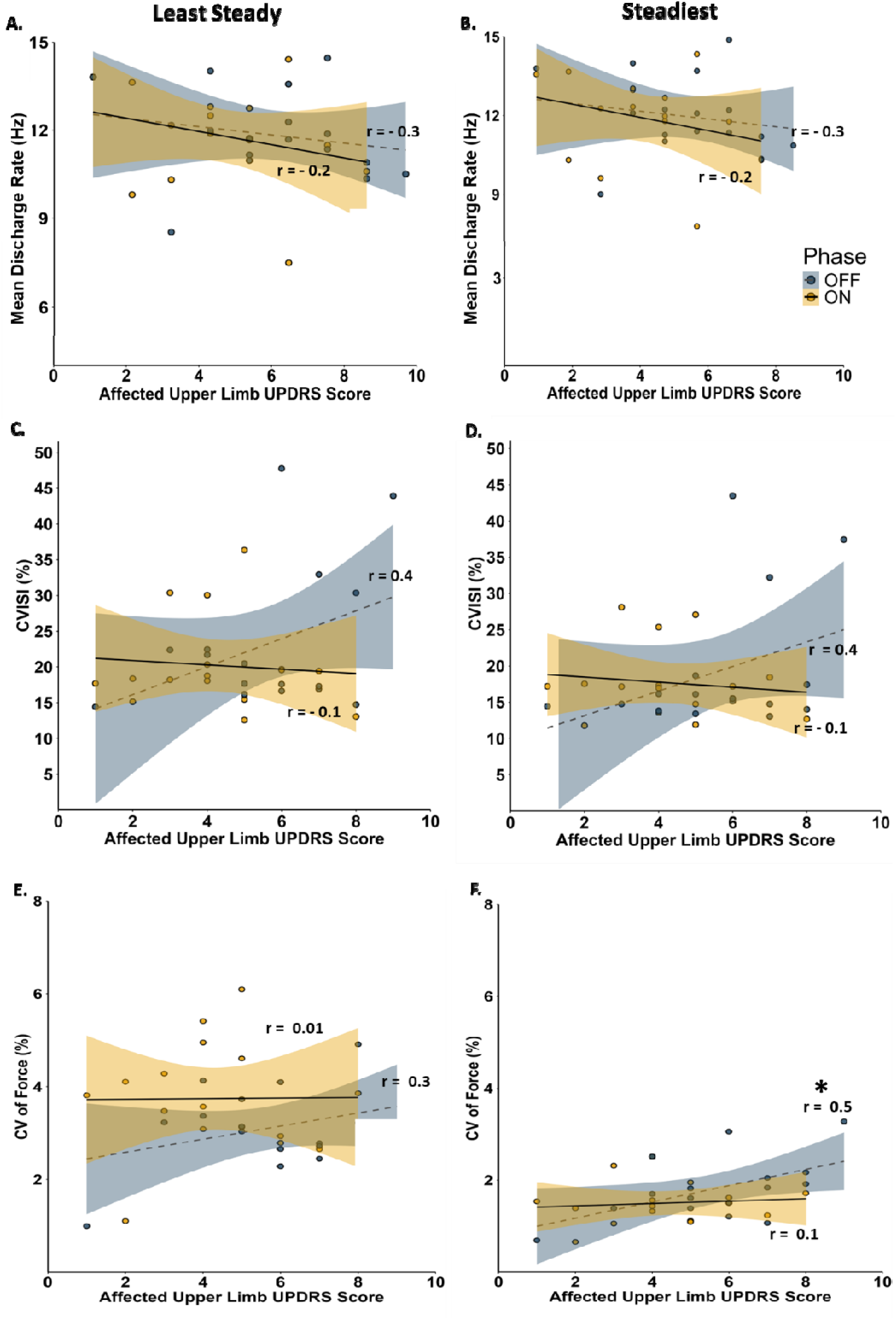
Relationship between affected upper limb UPDRS Score and (A) mean firing rate, (C) CVISI, and (E) CV of Force in the least steady period and (B) mean discharge rate, (D) CVISI, and (F) CV of Force in the steadiest period across all force levels. Each circle represents an individual participant-level mean, calculated by averaging across all MUs for that participant, during the ON (open circle) and OFF (filled circle) medication states.

### Motor Unit Coherence

No significant Medication state × Force interactions were observed in any frequency band (all p > 0.05; Table 2). A significant main effect of medication was observed in the Alpha (F(1, 371.2) = 6.9, p = 0.009; OFF: 0.52 ± 0.04; ON: 0.45 ± 0.04) and Tremor (F(1, 371.08) = 9.2, p = 0.003; OFF: 0.79 ± 0.10; ON: 0.65 ± 0.10) bands, with higher coherence during OFF compared with ON medication in both bands. Significant main effects of force were observed in the Beta (F(3, 371.9) = 3.47, p = 0.02), Delta (F(3, 371.1) = 5.15, p = 0.002), and Tremor (F(3, 371.1) = 3.42, p = 0.02) bands. Adjusted post-hoc comparisons showed that coherence was higher at 25% than at 5% MVC in the Beta band (p = 0.01). In the Delta band, coherence was higher at 10% than 5% MVC (p = 0.02) and 25% (p=0.02), and 15% was higher than 5% MVC (p = 0.04) and 25% MVC (p = 0.04). In the Tremor band, coherence was higher at 25% than 10% MVC (p = 0.02). Full coherence spectra of MU discharge patterns across different force levels and medication states are shown for all participants in Figure 8.

**Table 2.** Z-transformed coherence at each force level for each frequency band.

| Frequency Band | 5% |  | *p* | 10% |  | *p* | 15% |  | *p* | 25% |  | *p* |
| --- | --- | --- | --- | --- | --- | --- | --- | --- | --- | --- | --- | --- |
|  | OFF | ON |  | OFF | ON |  | OFF | ON |  | OFF | ON |  |
| **Alpha** | 0.48±0.32 | 0.42±0.22 | 1.0 | 0.54±0.37 | 0.45±0.23 | 1.0 | 0.57±0.58 | 0.41±0.29 | 0.4 | 0.46±0.31 | 0.49±0.27 | 1.0 |
| **Beta** | 0.27±0.12 | 0.27±0.11 | 1.0 | 0.30±0.14 | 0.30±0.10 | 1.0 | 0.33±0.11 | 0.29±0.15 | 1.0 | 0.30±0.15 | 0.36±0.19 | 0.4 |
| **Delta** | 2.03±1.63 | 2.07±1.39 | 1.0 | 2.68±1.40 | 2.37±1.04 | 1.0 | 2.53±1.40 | 2.31±1.32 | 1.0 | 2.00±1.17 | 2.10±1.01 | 1.0 |
| **Gamma** | 0.28±0.11 | 0.27±0.10 | 0.9 | 0.29±0.10 | 0.29±0.06 | 1.0 | 0.31±0.10 | 0.27±0.12 | 0.8 | 0.30±0.12 | 0.33±0.16 | 1.0 |
| **Tremor** | 0.79±0.74 | 0.69±0.53 | 1.0 | 0.90±0.74 | 0.70±0.48 | 0.8 | 0.84±0.73 | 0.63±0.49 | 0.3 | 0.61±0.50 | 0.57±0.35 | 1.0 |

Values are mean ± SD. Force is expressed as a percentage of maximum voluntary contraction (MVC). *p* values represent comparisons between the OFF and ON medication states at each force level.

**Figure 8.**
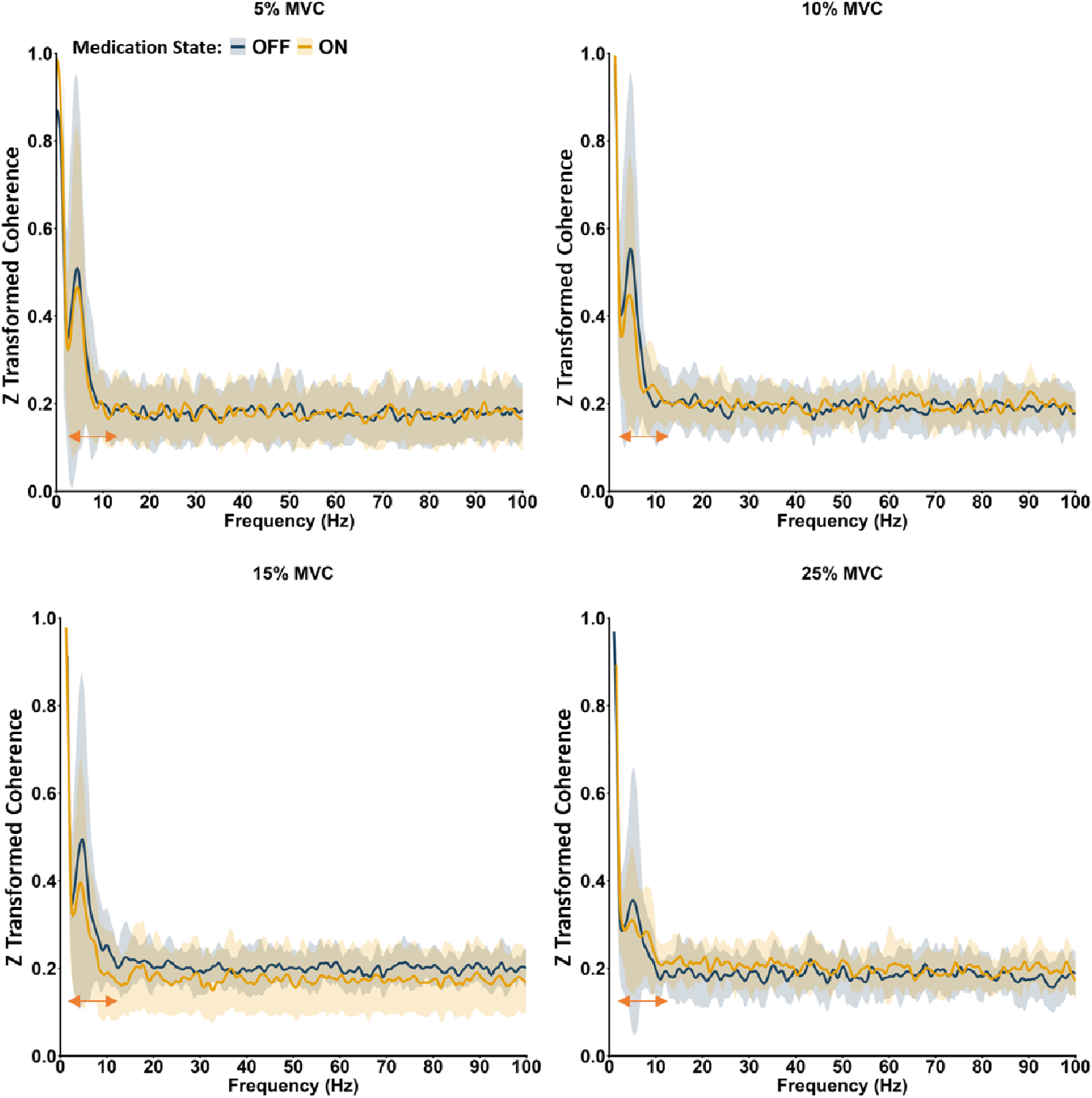
Full coherence spectra of motor unit discharge patterns across force levels. Lines represent the mean Fisher z-transformed coherence across participants for the OFF medication (blue) and ON medication (yellow) conditions at 5%, 10%, 15%, and 25% of maximal voluntary contraction (MVC). Shaded regions represent ±1 SD across participants at each frequency. Orange arrow illustrating the tremor frequency band (3-12 Hz). Hz, hertz; MVC, maxima voluntary contraction.

## Discussion

Comparing the steadiest and least steady contraction periods revealed that force control and individual MU discharge behavior vary within sustained contractions and are modified by dopaminergic medication in people with PD. Medication improved force steadiness during both contraction periods at lower force levels but not at 25% MVC. MU interspike interval variability was reduced more consistently during the least steady contraction period, whereas mean MU discharge rate did not differ between medication states or contraction periods. Plateau-wide analysis showed lower alpha- and tremor-band MU coherence during the ON state, indicating that medication also modified the frequency-specific organization of MU-pool activity across the contraction. Separating the two contraction periods showed that MU discharge variability and force control fluctuate within a sustained contraction and that medication had its most consistent effect on MU discharge variability during the period of poorer force control.

Force was consistently more variable during the least steady than the steadiest contraction period at every target force and in both medication states. Medication nevertheless improved force steadiness during both contraction periods at 5%, 10%, and 15% MVC, showing that its benefit extended across the within-contraction variation in force control rather than being confined to the least steady contraction period. The absence of an effect at 25% MVC may reflect a shift in the determinants of force fluctuations as contraction intensity increases. Greater coactivation of antagonist muscles at higher force levels may also contribute to force fluctuations, potentially reducing the relative influence of medication-related changes in individual MU behavior on force steadiness. Variability in individual MU discharges contributes substantially to force fluctuations at low forces (Laidlaw et al., 2000; Moritz et al., 2005; Taylor et al., 2003; Tracy et al., 2005), whereas common synaptic input to the MU pool becomes more prominent at moderate forces (Enoka & Farina, 2021; Negro et al., 2009; Tvrdy et al., 2025). Medication-related changes in individual MU behavior may therefore exert less influence on force steadiness as contraction intensity increases.

Dopaminergic medication had its most consistent effect of reducing MU variability during the least steady contraction period (∼12% across all forces). MU interspike interval variability decreased at every force during this period but was largely unchanged during the steadiest contraction period. Consequently, the greater MU interspike interval variability during the least steady than the steadiest contraction period in the OFF state was no longer evident in the ON state. This response corresponded with improved force steadiness in the ON state during the least steady contraction period at 5-15% MVC, but the patterns were not identical across force levels. MU interspike interval variability also decreased at 25% MVC without an improvement in force steadiness in the ON state, whereas steadiness improved during the steadiest contraction period at lower forces without corresponding reductions in MU interspike interval variability. Thus, medication-related improvements in force control were not limited to changes in individual MU discharge timing.

The location of the least steady contraction period further characterizes this within-contraction variation. It occurred in the second half of the plateau in 56% of OFF trials and 53% of ON trials and was therefore distributed approximately equally between the first and second halves and similarly across medication states. Medication improved the magnitude of force steadiness without altering when the least steady contraction period typically occurred within the 25-s plateau. Thus, force control fluctuated throughout the sustained contraction rather than declining progressively over time. Identifying both contraction periods capture variation that may be obscured by plateau-wide averaging or excluded when only the steadiest segment is analyzed.

Medication did not alter MUDR during either contraction period at any force. Thus, the improvement in force steadiness cannot be attributed to changes in average MU discharge rate. This result is consistent with evidence that biceps brachii MUDR does not differ between people with PD tested OFF medication and neurologically intact controls (Wilson et al., 2020), and with our previous observation that dopaminergic medication did not alter estimates of biceps brachii motoneuron excitability (Alaei et al., 2026). Earlier reports of reduced MUDR in PD primarily examined intrinsic hand muscles (Dietz et al., 1974; Milner-Brown et al., 1979), suggesting that the apparent difference may reflect muscle-specific control. Biceps brachii relies substantially on MU recruitment to increase force (Kukulka & Clamann, 1981), which may make its average discharge rate less sensitive to medication than that of distal muscles. The present findings therefore distinguish mean discharge rate from discharge regularity: medication changed CVISI despite leaving MUDR unchanged.

Recruitment thresholds were higher in the ON than OFF state at 10% and 25% MVC, whereas derecruitment thresholds did not change. Lower recruitment thresholds have previously been reported in the first dorsal interosseous of people with PD tested OFF medication relative to neurologically intact controls (Glendinning & Enoka, 1994). People with PD also exhibit greater agonist-antagonist coactivation (Busse et al., 2005; Keloth et al., 2021) and reduced reciprocal inhibition on the more affected side (Meunier et al., 2000). The higher recruitment thresholds in the ON state may therefore reflect medication-related changes in descending or spinal input, potentially including enhanced reciprocal inhibition and reduced antagonist coactivation. Because antagonist activity and reciprocal inhibition were not measured the mechanism cannot be determined from the present data. The positive association between recruitment and derecruitment thresholds in both medication states indicates that the relative ordering of MU activation and deactivation was broadly preserved.

Plateau-wide coherence provided complementary evidence that medication altered the organization of common synaptic input to the MU pool. Alpha- and tremor-band coherence were lower in the ON state, and these reductions did not depend on force level. By contrast, delta-band coherence, which overlaps the low-frequency neural drive most directly translated into force, did not differ between medication states. Muscle acts as a low-pass filter, so common input below approximately 5 Hz explains much of the variance in force, whereas higher-frequency inputs are attenuated (Farina & Negro, 2015; Negro et al., 2016). Reduced alpha- and tremor-band coherence should therefore not be interpreted as a reduction in the effective neural drive required to produce the target force. Instead, it may reflect less oscillatory synchronization in frequency ranges associated with PD tremor. When sufficiently strong, these oscillations can appear in motor output and contribute to force fluctuations (Negro et al., 2016). Because coherence was assessed across the entire plateau, these changes characterize overall MU-pool activity and cannot be attributed to either contraction period.

Clinical motor severity was related to force control primarily during the OFF state. Higher affected-limb MDS-UPDRS III scores were associated with greater force variability during the steadiest contraction period; the relationship during the least steady contraction period was also positive but not significant. MU interspike interval variability showed moderate positive, although nonsignificant, relationships with clinical severity during both contraction periods in the OFF state. Mean MU discharge rate was unrelated to clinical severity in either medication state. During the ON state, neither force variability nor MU interspike interval variability was associated with MDS-UPDRS III scores during either contraction period. Thus, relationships between motor symptom severity and variability in force output and MU discharge timing were more evident in the OFF state and largely absent in the ON state. Previous studies reported positive relationships between affected upper limb UPDRS III scores and mean MU discharge rate (Nishikawa et al., 2021, 2025), but their cohorts had shorter disease durations (∼ 3 years), and both studies examined the lower-limb vastus lateralis muscle during ON medication. Overall, this pattern indicates that dopaminergic medication weakened the relationship between motor symptom severity and variability in MU discharge timing and force output, consistent with reduced expression of clinical motor deficits in these measures during the ON state. A limitation of the present study is that the OFF condition was a defined OFF state of a 12-h medication-withdrawal period. Although this approach balances medication withdrawal with participant feasibility, it may not fully exclude longer-duration effects of dopaminergic medication. Thus, ON–OFF differences should be interpreted as differences between medication states rather than as free of residual medication effects. In six of 16 participants the ON-state assessment, due to patient feasibility, followed an afternoon rather than morning dose. This variability may have influenced the magnitude or timing of the dopaminergic medication response.

In summary, dopaminergic medication improved low-force steadiness in people with PD while reducing MU discharge-timing variability, particularly during the least steady contraction period. Mean MU discharge rate was unchanged, whereas recruitment threshold and plateau-wide alpha- and tremor-band coherence showed selective medication effects. These results suggest that dopaminergic medication improves force regulation by modifying the timing and frequency-specific coordination of MU activity rather than uniformly increasing MU discharge rate. Comparing the steadiest and least steady contraction periods reveals within-contraction changes that are not represented by conventional plateau-wide analyses and provides a useful framework for studying impaired force control in PD.

## Data Availability Statement

The data supporting the conclusions of this article will be made available by the authors, upon reasonable request without undue reservation.

## Ethics Statement

This study was approved by the Clinical Research Ethics Board at the University of British Columbia and the Interior Health Authority through a Harmonized review process (H22-02896). All procedures were conducted in accordance with the Declaration of Helsinki, local health authority regulations, and institutional guidelines. Written informed consent was obtained from all participants prior to their inclusion in this study.

## Author Contributions

P.A: Data collection, methodology, data curation and analysis, writing original draft, and revising. A.H: Methodology, review & editing document. D.J.W: Conceptualization, methodology, clinical resources, review & editing document. J.M.J: Supervision, conceptualization, methodology, experimental resources, writing & editing document. All authors have read and agreed to the published version of the manuscript.

## Acknowledgments

We sincerely thank all the participants for their time and dedication, which were vital to the success of this research. We also extend our gratitude to the Kelowna and Vernon Parkinson’s Disease Community Groups for their valuable assistance with participant recruitment. We also gratefully acknowledge Gregory E.P. Pearcey for his help, guidance, and support throughout this research.

