## Supplemental Materials for "Dopaminergic Medication Improves Force Control and Modifies Motor Unit Activity During the Steadiest and Least Steady Periods in Parkinson’s Disease Force control and motor units in Parkinson’s Disease"

***Supplemental Material***

***Table1.*** Recruitment and Derecruitment Thresholds During ON and OFF Phases (Mean ± SD)

| **Force Level** | **Recruitment Threshold (%)** | | ***p*** | **Derecruitment Threshold (%)** | | ***p*** | **Recruitment vs Derecruitment**  **Threshold** | |
| --- | --- | --- | --- | --- | --- | --- | --- | --- |
|  | **OFF** | **ON** |  | **OFF** | **ON** |  | ***p* OFF** | ***p* ON** |
| 5% | 1.0 ± 1.2 | 1.0 ± 1.3 | 0.6 | 1.5 ± 1.4 | 1.4 ± 1.6 | 0.9 | < 0.001* | < 0.001* |
| 10% | 2.5 ± 2.9 | 2.9 ± 2.7 | 0.02* | 3.6 ± 3.0 | 2.8 ± 2.9 | 0.2 | < 0.001* | 1.0 |
| 15% | 5.4 ± 4.0 | 4.5 ± 3.9 | 0.1 | 5.5 ± 4.2 | 5.2 ± 4.1 | 1.0 | 0.2 | 1.0 |
| 25% | 8.9 ± 6.8 | 10.7 ± 6.4 | 0.02* | 9.8 ± 6.7 | 10.7 ± 7.0 | 0.4 | 1.0 | 0.5 |
| *OFF, OFF phase; ON, ON phase. p, p-value for comparison between ON and OFF phases. p OFF, p-value for the comparison between recruitment and derecruitment thresholds within the OFF phase. p ON, p-value for the comparison between recruitment and derecruitment thresholds within the ON phase*. **, p < 0.001.* | | | | | | | | |

***Table 2****.* Cohen’s d effect sizes for pairwise comparisons of motor unit recruitment and derecruitment threshold across medication and steadiness conditions

| **Force** | **Contrast** | **Cohen’s *d*** |
| --- | --- | --- |
| **5%** | ON Recruitment threshold − OFF Recruitment threshold | 0.13 |
|  | ON Derecruitment threshold − OFF Derecruitment threshold | 0.08 |
|  | OFF Recruitment threshold − OFF Derecruitment threshold | 0.41 |
|  | ON Recruitment threshold − ON Derecruitment threshold | 0.46 |
|  | ON Recruitment threshold − OFF Derecruitment threshold | 0.54 |
| **10%** | ON Recruitment threshold − OFF Recruitment threshold | 0.29 |
|  | ON Derecruitment threshold − OFF Derecruitment threshold | 0.20 |
|  | OFF Recruitment threshold − OFF Derecruitment threshold | 0.49 |
|  | ON Recruitment threshold − ON Derecruitment threshold | 0.00 |
|  | ON Recruitment threshold − OFF Derecruitment threshold | 0.20 |
| **15%** | ON Recruitment threshold − OFF Recruitment threshold | 0.26 |
|  | ON Derecruitment threshold − OFF Derecruitment threshold | 0.08 |
|  | OFF Recruitment threshold − OFF Derecruitment threshold | 0.03 |
|  | ON Recruitment threshold − ON Derecruitment threshold | 0.25 |
|  | ON Recruitment threshold − OFF Derecruitment threshold | 0.33 |
| **25%** | ON Recruitment threshold − OFF Recruitment threshold | 0.31 |
|  | ON Derecruitment threshold − OFF Derecruitment threshold | 0.08 |
|  | OFF Recruitment threshold − OFF Derecruitment threshold | 0.20 |
|  | ON Recruitment threshold − ON Derecruitment threshold | 0.03 |
|  | ON Recruitment threshold − OFF Derecruitment threshold | 0.11 |

*OFF, OFF phase. ON, ON phase*

***Table 3***. Estimated marginal means, standard errors, and 95% confidence limits for motor unit recruitment and derecruitment threshold across medication and steadiness conditions

| **Force** | **Factor** | **Emmean** | **SE** | **95% CI** |
| --- | --- | --- | --- | --- |
| **5%** | Recruitment threshold OFF | 1.02 | 0.21 | [−1.47, −0.56] |
|  | Recruitment threshold ON | −1.24 | 0.22 | [−1.69, −0.79] |
|  | Derecruitment threshold OFF | −0.31 | 0.20 | [−0.77, 0.14] |
|  | Derecruitment threshold ON | −0.46 | 0.22 | [−0.91, −0.01] |
| **10%** | Recruitment threshold OFF | −0.24 | 0.22 | [−0.69, 0.20] |
|  | Recruitment threshold ON | 0.25 | 0.21 | [−0.18, 0.69] |
|  | Derecruitment threshold OFF | 0.59 | 0.22 | [0.15, 1.04] |
|  | Derecruitment threshold ON | 0.25 | 0.21 | [−0.18, 0.69] |
| **15%** | Recruitment threshold OFF | 1.05 | 0.20 | [0.64, 1.47] |
|  | Recruitment threshold ON | 0.43 | 0.21 | [0.01, 0.86] |
|  | Derecruitment threshold OFF | 1.00 | 0.20 | [0.58, 1.42] |
|  | Derecruitment threshold ON | 0.86 | 0.21 | [0.44, 1.29] |
| **25%** | Recruitment threshold OFF | 1.40 | 0.21 | [0.96, 1.83] |
|  | Recruitment threshold ON | 1.86 | 0.22 | [1.43, 2.30] |
|  | Derecruitment threshold OFF | 1.70 | 0.21 | [1.26, 2.13] |
|  | Derecruitment threshold ON | 1.81 | 0.21 | [1.38, 2.25] |

*OFF, OFF phase. ON, ON phase. Emmean = estimated marginal mean; SE = standard error; 95% CI = 95% confidence interval.*

**Table 4**. Motor unit discharge rate and CVISI during ON and OFF medication phases in the least steady and steadiest period of a contraction (Mean ± SD)

| **Motor Unit Discharge Rate (Hz)** | | | | | | | | | |
| --- | --- | --- | --- | --- | --- | --- | --- | --- | --- |
| **Force Level** | **Least Steady** | | ***p*** | **Steadiest** | | ***P*** | ***p* OFF** | ***p* ON** | ***p Optimal*** |
|  | **OFF** | **ON** |  | **OFF** | **ON** |  |  |  |  |
| **5%** | 11.4 ± 2.2 | 11.1 ± 2.3 | 0.2 | 11.7 ± 2.1 | 11.3 ± 1.9 | 0.1 | 0.3 | 0.4 | 1.0 |
| **10%** | 11.6 ± 2.7 | 12.1 ± 2.7 | 0.5 | 11.9 ± 2.9 | 12.3 ± 2.5 | 0.5 | 0.7 | 0.6 | 0.06 |
| **15%** | 12.4 ± 2.9 | 12.5 ± 2.5 | 1.0 | 12.5 ± 2.9 | 12.5 ± 2.7 | 0.9 | 0.9 | 1.0 | 1.0 |
| **25%** | 13.2 ± 3.2 | 13.3 ± 2.7 | 0.6 | 13.0 ± 2.9 | 13.3 ± 2.9 | 0.9 | 1.0 | 0.9 | 0.9 |
| **CVISI (%)** | | | | | | | | | |
|  | **Least Steady** | | ***p*** | **Steadiest** | | ***p*** | ***p* OFF** | ***p* ON** |  |
|  | **OFF** | **ON** |  | **OFF** | **ON** |  |  |  |  |
| **5%** | 23.5 ± 16.3 | 21.7 ± 16.9 | 0.04 * | 19.2 ± 13.1 | 17.8 ± 10.0 | 1.0 | < 0.001 * | 0. 1 | < 0.001 * |
| **10%** | 23.0 ± 15.4 | 18.6 ± 12.7 | 0.002 * | 18.5 ± 14.1 | 16.3 ± 9.4 | 0.8 | < 0.001 * | 0.1 | < 0.001 * |
| **15%** | 23.1 ± 15.8 | 18.0 ± 9.3 | 0.002 * | 19.8 ± 11.7 | 16.4 ± 10.2 | 0.03 * | 0.002 * | 0.06 | < 0.001 * |
| **25%** | 22.1 ± 15.3 | 18.4 ± 10.6 | 0.02 * | 18.1 ± 11.3 | 16.7 ± 8.6 | 0.7 | 0.003 * | 0.3 | < 0.001 * |
| *Motor unit discharge rate and CVISI at 5, 10, 15 and 25% MVC during ON and OFF medication phases. OFF, OFF phase. ON, ON phase. P, p-value for comparison between ON and OFF phases. P OFF, p-value for the comparison between the least steady and steadiest periods within the OFF phase. P ON, p-value for the comparison between least steady and steadiest periods within the ON phase*. *Hz, Hertz*. **, p < 0.05* | | | | | | | | | |

***Table 5****.* Cohen’s d effect sizes for pairwise comparisons of motor unit discharge rate and CVISI across medication and steadiness conditions

| **Force** | **Contrast** | **Cohen’s *d*** | **Cohen’s *d*** |
| --- | --- | --- | --- |
|  |  | **MUDR** | **CVISI** |
| **5%** | ON Least steady − OFF Least steady | 0.19 | 0.28 |
|  | ON Steady − OFF Steady | 0.26 | 0.04 |
|  | OFF Least steady − OFF Steady | 0.17 | 0.49 |
|  | ON Least steady − ON Steady | 0.11 | 0.25 |
|  | OFF Least steady − ON Steady | 0.37 | 0.53 |
| **10%** | ON Least steady − OFF Least steady | 0.16 | 0.36 |
|  | ON Steady − OFF Steady | 0.11 | 0.09 |
|  | OFF Least steady − OFF Steady | 0.15 | 0.47 |
|  | ON Least steady − ON Steady | 0.10 | 0.21 |
|  | OFF Least steady − ON Steady | 0.01 | 0.57 |
| **15%** | ON Least steady − OFF Least steady | 0.04 | 0.37 |
|  | ON Steady − OFF Steady | 0.08 | 0.29 |
|  | OFF Least steady − OFF Steady | 0.05 | 0.35 |
|  | ON Least steady − ON Steady | 0.02 | 0.27 |
|  | OFF Least steady − ON Steady | 0.09 | 0.64 |
| **25%** | ON Least steady − OFF Least steady | 0.09 | 0.31 |
|  | ON Steady − OFF Steady | 0.06 | 0.12 |
|  | OFF Least steady − OFF Steady | 0.02 | 0.38 |
|  | ON Least steady − ON Steady | 0.05 | 0.18 |
|  | OFF Least steady − ON Steady | 0.11 | 0.49 |

*OFF, OFF phase. ON, ON phase. MUDR, motor unit discharge rate; CVISI, coefficient of variation of interspike interval.*

***Table 6***. Estimated marginal means, standard errors, and 95% confidence limits for motor unit discharge rate and CVISI across medication and steadiness conditions.

|  |  | **MUDR** | | | | **CVISI** | | |
| --- | --- | --- | --- | --- | --- | --- | --- | --- |
| **Force** | **Condition** | **Emmean** | **SE** | **95% CI** | **Emmean** | | **SE** | **95% CI** |
| **5%** | OFF Least | 11.20 | 0.31 | [10.55, 11.85] | 3.05 | | 0.12 | [2.80, 3.29] |
|  | ON Least | 10.83 | 0.29 | [10.19, 11.48] | 2.94 | | 0.11 | [2.69, 3.18] |
|  | OFF Steady | 11.54 | 0.31 | [10.89, 12.18] | 2.85 | | 0.11 | [2.61, 3.10] |
|  | ON Steady | 11.04 | 0.32 | [10.40, 11.69] | 2.84 | | 0.10 | [2.59, 3.08] |
| **10%** | OFF Least | 11.40 | 0.45 | [10.46, 12.34] | 2.94 | | 0.09 | [2.75, 3.13] |
|  | ON Least | 11.75 | 0.45 | [10.81, 12.69] | 2.78 | | 0.089 | [2.60, 2.97] |
|  | OFF Steady | 11.74 | 0.44 | [10.79, 12.68] | 2.74 | | 0.09 | [2.55, 2.93] |
|  | ON Steady | 11.98 | 0.43 | [11.04, 12.92] | 2.70 | | 0.08 | [2.51, 2.88] |
| **15%** | OFF Least | 12.10 | 0.43 | [11.15, 13.04] | 2.92 | | 0.08 | [2.73, 3.10] |
|  | ON Least | 12.00 | 0.45 | [11.05, 12.95] | 2.76 | | 0.09 | [2.58, 2.95] |
|  | OFF Steady | 12.21 | 0.45 | [11.26, 13.16] | 2.77 | | 0.08 | [2.59, 2.96] |
|  | ON Steady | 12.04 | 0.46 | [11.08, 12.99] | 2.66 | | 0.08 | [2.47, 2.84] |
| **25%** | OFF Least | 12.81 | 0.51 | [11.73, 13.89] | 2.93 | | 0.07 | [2.78, 3.07] |
|  | ON Least | 13.02 | 0.52 | [11.94, 14.11] | 2.79 | | 0.07 | [2.64, 2.94] |
|  | OFF Steady | 12.77 | 0.52 | [11.68, 13.86] | 2.76 | | 0.06 | [2.61, 2.91] |
|  | ON Steady | 12.90 | 0.52 | [11.82, 13.99] | 2.71 | | 0.06 | [2.56, 2.86] |

*OFF, OFF phase. ON, ON phase. Emmean = estimated marginal mean; SE = standard error; 95% CI = 95% confidence interval. MUDR, motor unit discharge rate; CVISI, coefficient of variation of interspike interval*

***Table 7****.* Cohen’s d effect sizes for pairwise comparisons of force steadiness across medication and steadiness conditions.

| **Force** | **Contrast** | **Cohen’s *d*** |
| --- | --- | --- |
| **5%** | ON Least steady − OFF Least steady | 0.61 |
|  | ON Steady − OFF Steady | 0.61 |
|  | OFF Least steady − OFF Steady | 1.58 |
|  | ON Least steady − ON Steady | 1.59 |
|  | OFF Least steady − ON Steady | 2.20 |
| **10%** | ON Least steady − OFF Least steady | 0.54 |
|  | ON Steady − OFF Steady | 0.55 |
|  | OFF Least steady − OFF Steady | 1.97 |
|  | ON Least steady − ON Steady | 1.97 |
|  | OFF Least steady − ON Steady | 2.51 |
| **15%** | ON Least steady − OFF Least steady | 0.97 |
|  | ON Steady − OFF Steady | 0.61 |
|  | OFF Least steady − OFF Steady | 2.20 |
|  | ON Least steady − ON Steady | 1.84 |
|  | OFF Least steady − ON Steady | 2.81 |
| **25%** | ON Least steady − OFF Least steady | 0.39 |
|  | ON Steady − OFF Steady | 0.34 |
|  | OFF Least steady − OFF Steady | 1.93 |
|  | ON Least steady − ON Steady | 1.88 |
|  | OFF Least steady − ON Steady | 2.27 |

*OFF, OFF phase. ON, ON phase*

***Table 8***. Estimated marginal means, standard errors, and 95% confidence limits for force steadiness across medication and steadiness conditions.

| **Force** | **Condition** | **Emmean** | **SE** | **95% CI** |
| --- | --- | --- | --- | --- |
| **5%** | OFF Least | 1.65 | 0.13 | 1.38–1.92 |
|  | ON Least | 1.39 | 0.14 | 1.12–1.66 |
|  | OFF Steady | 0.96 | 0.13 | 0.69–1.23 |
|  | ON Steady | 0.70 | 0.12 | 0.43–0.97 |
| **10%** | OFF Least | 1.19 | 0.10 | 0.97–1.40 |
|  | ON Least | 0.99 | 0.11 | 0.78–1.20 |
|  | OFF Steady | 0.48 | 0.10 | 0.27–0.69 |
|  | ON Steady | 0.28 | 0.12 | 0.07–0.49 |
| **15%** | OFF Least | 1.06 | 0.08 | 0.89–1.22 |
|  | ON Least | 0.72 | 0.09 | 0.53–0.89 |
|  | OFF Steady | 0.31 | 0.08 | 0.14–0.47 |
|  | ON Steady | 0.10 | 0.08 | −0.06–0.26 |
| **25%** | OFF Least | 0.87 | 0.08 | 0.68–1.05 |
|  | ON Least | 0.72 | 0.09 | 0.53–0.91 |
|  | OFF Steady | 0.14 | 0.09 | −0.04–0.33 |
|  | ON Steady | 0.02 | 0.09 | −0.17–0.21 |

*OFF, OFF phase. ON, ON phase. Emmean = estimated marginal mean; SE = standard error; 95% CI = 95% confidence interval.*
